# On-site eDNA detection of species using ultra-rapid mobile PCR

**DOI:** 10.1101/2020.09.28.314625

**Authors:** Hideyuki Doi, Takeshi Watanabe, Naofumi Nishizawa, Tatsuya Saito, Hisao Nagata, Yuichi Kameda, Nobutaka Maki, Kousuke Ikeda, Takashi Fukuzawa

**Author notes:** Corresponding author: Hideyuki Doi. Author Contributions HD, TW, NN, and TF designed and developed this study and protocol. TW, NN, TS, HN, YK, NM, and KI contributed to measurement. HD analyzed the data and wrote the initial draft of the manuscript, and all authors critically reviewed the manuscript.

## Abstract

Molecular methods, including environmental DNA (eDNA) methods, provide essential information for biological and conservation sciences. Molecular measurements are often performed in the laboratory, which limits their scope, especially for rapid on-site analysis. eDNA methods for species detection provide essential information for the management and conservation of species and communities in various environments. We developed an innovative novel method for on-site eDNA measurements using an ultra-rapid mobile PCR platform. We tested the ability of our method to detect the distribution of silver carp, Hypophthalmichthys molitrix, an invasive fish in Japanese rivers and lakes. Our method reduced the measurement time to 30 min and provided high detectability of aquatic organisms compared to the national observation surveys using multiple fishing nets and laboratory extraction/detection using a benchtop qPCR platform. Our on-site eDNA method can be immediately applied to various taxa and environments.

## Introduction

Molecular technologies, such as species identification and gene expression analyses, provide essential information for biological and conservation sciences. However, even with advances in techniques in the last decades, molecular measurements in the laboratory may take a day or more. Ultra-rapid methods from DNA collection to detection are still not well developed (1), especially for environmental DNA (eDNA) analysis, which uses water or soil samples to track the presence of target species (2, 3).

eDNA analysis is a useful method to investigate the distribution of aquatic and terrestrial organisms (4–6). Approaches using eDNA have provided essential information for ecological management and conservation, facilitating the detection of various kinds of organisms, including endemic, invasive, or parasitic species (2, 6, 7).

eDNA measurements have been mainly performed by quantitative real-time PCR (qPCR, 4–7). However, it is limited to laboratory analysis and laboratory processing can take many hours. These time delays often limit the range of uses for on-site eDNA detection (9, 10). Field-portable DNA extraction and PCR platforms offer the potential to change species detection by eDNA on site (8–10). However, these approaches still take a similar time to laboratory measurements.

Here, we developed a new innovative method for the field processing of eDNA samples and measurements using an ultra-rapid mobile PCR platform (hereafter, mobile PCR) to reduce the measurement time to 30 min and maintain high detectability of aquatic organisms. We demonstrated its on-site use to detect the distribution of silver carp, Hypophthalmichthys molitrix, an invasive fish in Japanese rivers and lakes. We compared the on-site eDNA measurement to the laboratory extraction and detection using a benchtop qPCR platform and the national survey to confirm the performance.

## Results

We detected the eDNA of H. molitrix by on-site measurement at 11 out of 15 sites (Fig. 1), including almost all sites (except a site) where the distribution was recorded by a national survey using multiple fishing nets. The eDNA and the national surveys were significantly matched (Cohen's Kappa tests; Survey 1: Kappa = 0.865, p = 0.00072, Survey 2: Kappa = 1.00, p = 0.0027). We took approximately 30 min to carry out all on-site eDNA extraction and measurements.

**Figure 1.**
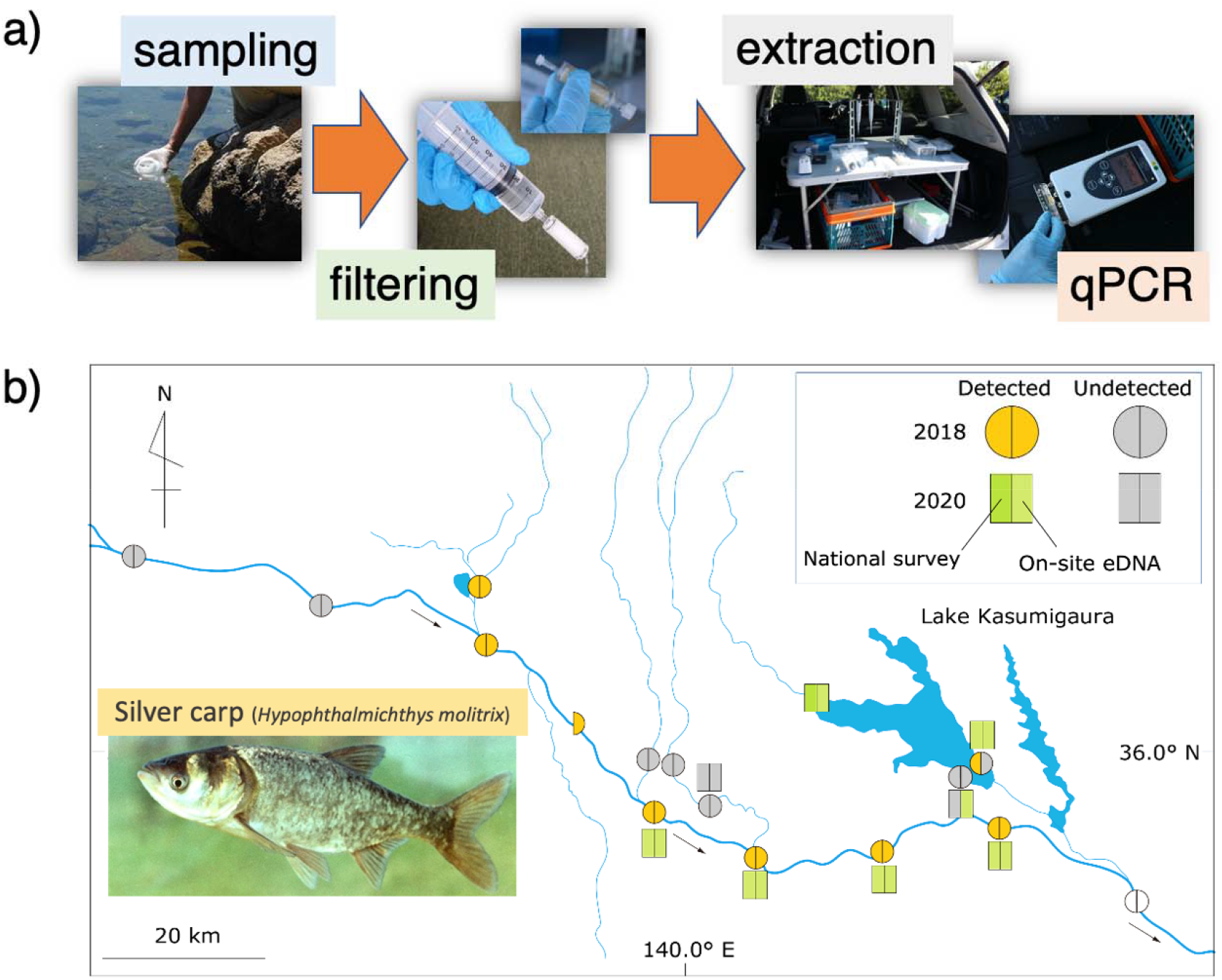
a) Illustration of the on-site eDNA sampling and measurement by mobile PCR and b) the map for detection of Hypophthalmichthys molitrix eDNA detection by Survey 1 (2018) and 2 (2020) using mobile PCR and H. molitrix observations from the national survey using multiple fishing nets.

In Survey 2, we also detected the eDNA by the laboratory methods using benchtop qPCR at all sites detected by our on-site method (Fig. 1). The relationship between the cycle timing (Ct) of the mobile PCR and eDNA concentration (or Ct) of qPCR was significant (Fig. 2a, b, LM, p < 0.001). The Ct of mobile PCR was larger than that of qPCR, because the DNA concentration in the field-extracted samples was lower.

**Figure 2.**
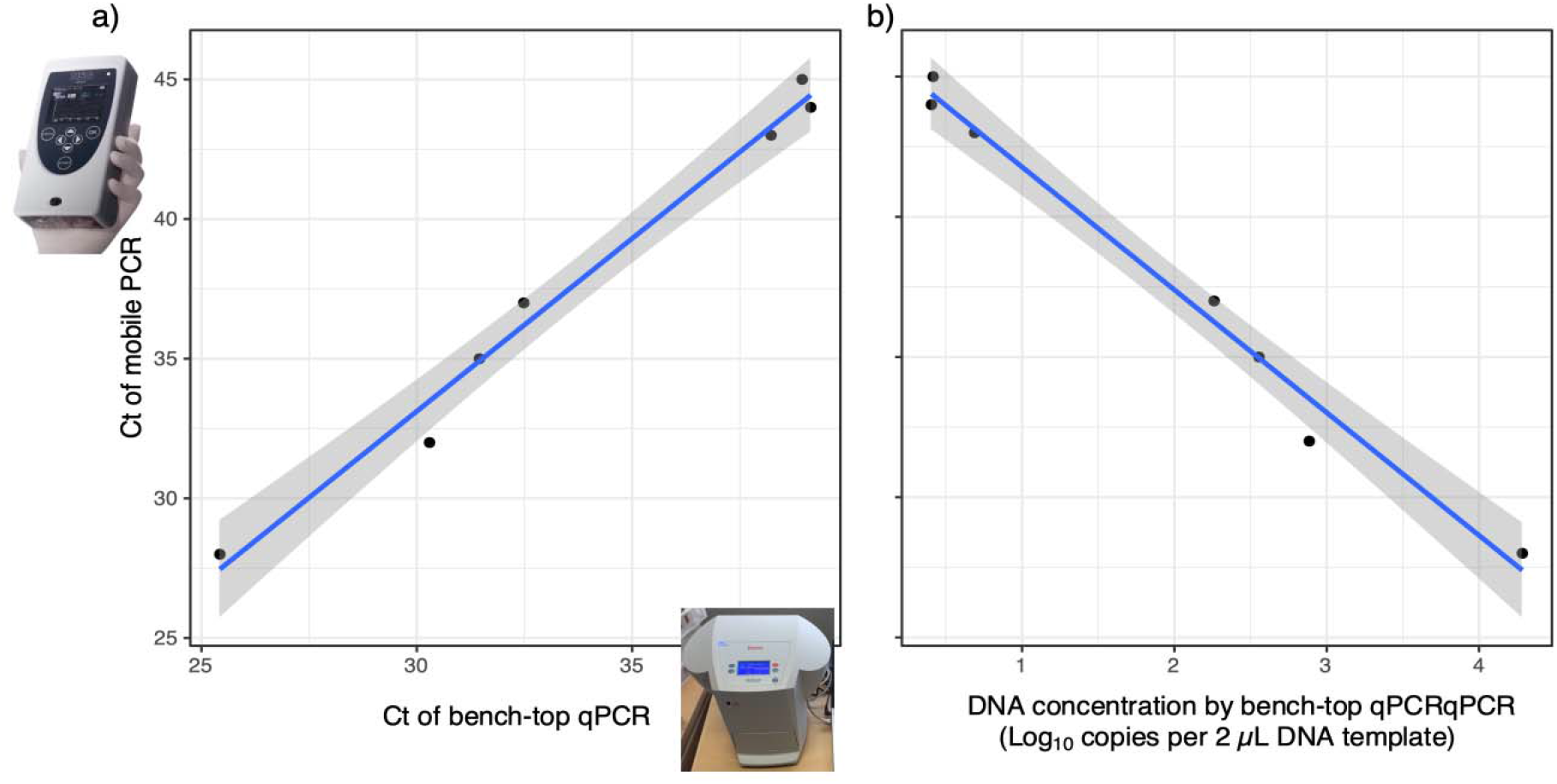
Relationships for on-site extracted DNA measurements between a) cycle timing (Ct) of qPCR and Ct of mobile PCR, b) eDNA concentration by qPCR and Ct of mobile PCR. The line and gray area indicate the GLM regression and 95% CI, respectively. All LMs were significant: a) p= 0.0000012, r^2^ =0.9802, b) p= 0.0000031, r^2^ = 0.9711).

## Discussion

Our ultra-rapid on-site eDNA extraction and measurement method using mobile PCR successfully detected the eDNA of H. molitrix, and analysis took only 30 min. Our method can be applied to many other taxa, including viruses and bacteria and to vertebrates using specific primers. We sampled water from aquatic ecosystems, but the method can be applied to terrestrial systems. For example, Valentin et al. (11) evaluated terrestrial insects on forest leaves by spraying and collecting water. The mobile PCR platform can also perform multiplex PCR for a few independent DNA measurements. Using multiplex PCR, we can detect species co-existence, for example, for close host-parasites interactions. Therefore, our method has high potential for use with various taxa in different environments, including terrestrial and marine ecosystems.

This ultra-rapid methods can immediately be applied to broad science fields, such as human health (12) and food science (13). For example, Medema et al. (12) detected SARS-CoV-2 RNA from wastewater to evaluate the spread of COVID-19. Our method can be applied to detect RNA viruses, such as SARS-CoV-2, using reverse-transcription qPCR.

## Materials and Methods

### Study sites

We conducted field surveys in the Tone River and Lake Kasumigaura: Survey 1: on-site detection only and Survey 2: on-site detection and laboratory measurement. We conducted field surveys twice on 24 July and 4 August 2018 and 17–18 June 2020 for Survey 1 and 2. We performed sampling and eDNA measurement at 15 sites at 9 of the 15 sites for Survey 1 and 2, respectively.

### Water sampling and filtration

We collected one 500-mL sample per site using a bleached bucket. We filtered the water using a 0.45-μm Sterivex cartridge filter (Sterivex, Merck Millipore, Burlington, MA, USA) using a 50-mL syringe.

For Survey 2, we collected the water and filtered it through a 0.45-μm Sterivex twice. Then, 1.6 mL of RNAlater (Sigma-Aldrich, St. Louis, MO, USA) was added into a filtered 0.45-μm Sterivex (15). The all equipment was bleached with 10% commercial bleach (ca. 0.6% sodium hypochlorite) and washed with DNA-free distilled water (DW).

### On-site DNA extraction from Sterivex

We used the Kaneka Simple DNA Extraction Kit v.2 (Kaneka, Tokyo, Japan) for on-site DNA extraction from Sterivex. We injected 500 μL of Solution A of the kit into the Sterivex and shook it by hand for 1 min. We collected the Solution A buffer in the Sterivex by connecting the syringe with the connecting tube. We added 70 μL of Solution B into the collected Solution A buffer in a 1.5-ml microtube and centrifuged (~2000 g) with a portable centrifuge for 3 min. We collected 200 μL of the supernatant solution.

### On-site DNA measurement using mobile PCR

We used a primer and probe set to detect H. molitrix (14). We checked the primer specificity for other related species such as carp in Japan using NCBI Primer-BLAST (https://www.ncbi.nlm.nih.gov/tools/primer-blast/index.cgi) and confirmed the specificity for Japanese carp species.

A day before sampling, we made a PCR pre-mix with preliminary mixing of the master mix and primer-probe to bring it on site. Each TaqMan reaction contained 900 nM of each primer (forward and reverse), 400 nM TaqMan-Probe, 0.1-U/μL qPCR master mix (KAPA3G Plant PCR Kit; Millipore-Sigma, Darmstadt, Germany). The final volume for PCR pre-mix was made up to 14.4 μL by adding DW. The PCR pre-mix was stored in a cooler until PCR measurement.

Immediately after on-site DNA extraction, we measured the eDNA using PicoGene PCR1100 (mobile qPCR; Nippon Sheet Glass, Sagamihara, Japan). We added 1.6 μL of the eDNA solution in the tube of the pre-mix. We centrifuged the tube and then injected 14.4 μL of 16 μL into the flow path of PicoGene PCR1100 to reduce air bubbles.

The PCR conditions were as follows: 95 °C for 15 s, followed by 50 cycles of 95 °C for 3.5 s, and 62 °C for 10 s. In the laboratory, we performed a no-template control (NTC) using DW after the mixture preparation as a regents control. We performed an NTC using DW after all PCR measurements in the day (PCR control).

### Laboratory DNA measurement

For Survey 2, we extracted the DNA from the RNAlater-fixed Sterivex filters and purified using the DNeasy Blood and Tissue Kit (Qiagen, Hilden, Germany) according to Miya et al. (15).

We quantified the eDNA using the PikoReal Real-Time PCR (Thermo Fisher Scientific, Waltham, MA, USA). In the laboratory qPCR, we used the same primer-probe set of on-site measurements and the PCR template mix as in our previous studies (4). Each TaqMan reaction contained 900 nM of each primer (forward and reverse), 125 nM TaqMan-Probe, 5-μL qPCR master mix (TaqPath; Thermo Fisher Scientific), and 1.0 μL of the eDNA solution. The final volume for PCR was 10 μL by adding DW. The concentration of eDNA solution (10% of the PCR template) is the same as the mobile qPCR. The qPCR conditions were as follows: 50 °C for two min, 95 °C for 10 min, followed by 50 cycles of 95 °C for 15 s, and 60 °C for 60 s. We performed four replicates for each sample and NTC (N = 4).

We used a dilution series of 10000, 1000, 100, and 10 copies per PCR reaction (N = 4) for the standard curve using the target DNA cloned into a plasmid. The R^2^ values of the standard curves ranged from 0.989 to 0.994 (PCR efficiencies = 93.1–102.0%). We did not detect any positives from the controls for mobile PCR and qPCR, and confirmed no cross-contamination in all eDNA measurements.

### Limit of detection (LOD) test

We performed an LOD test for both mobile and qPCR as per the above PCR conditions. We used 1, 2, 4, and 8 copies of the positive control per PCR template with four replicates and detected two copy of the positive control (1/4 replicates). Thus, we determined that the LOD was two copies for both mobile and qPCR.

### Distribution data

We obtained a capture survey dataset from the Ministry of Land, Infrastructure, Transport and Tourism, Japan (http://www.nilim.go.jp/lab/fbg/ksnkankyo/index.html). The national fish survey was conducted using multiple fishing gears (casting, gill, and shin net) in 2014. The survey was conducted in three seasons, including spring, summer, and autumn, and H. molitrix was observed in multiple seasons.

### Statistical analysis

All statistical analyses were conducted using R ver. 4.0.2.We calculated Cohen’s Kappa value to compare the detection probability of H. molitrix distribution between eDNA and the national surveys with the R "irr" package ver. 0.84, with equally weighted data. To test the regression between eDNA concentration estimated by qPCR and the Ct of mobile PCR, we performed linear models (LMs) using “lm” function.

## Data availability

All data are available in the Supplementary Table S1.

## Acknowledgments

We thank Teruhiko Takahara for his helpful comments on our manuscript. This study was supported by the Environment Research and Technology Development Fund (4-1602, 4-2004).

## Notes

Competing Interest Statement: There are no conflicts of interest to declare.

### Competing Interest Statement

The authors have declared no competing interest.

